# Positive feedback loop of regulating ERK phosphorylation in mESCs mediated by Etv5-Tet2-Fgfr2 axis

**DOI:** 10.1101/560334

**Authors:** Chen Fan, Kui Zhu, Yuan Liu, Mengyao Zhang, Hongxia Cao, Na Li, Yan Wang, Jinlian Hua, Huayan Wang, Shiqiang Zhang

**Affiliations:** College of Veterinary Medicine, Shaanxi Center of Stem Cells Engineering & Technology, Northwest A&F University; College of Innovation and Experiment, Northwest A&F University

**Keywords:** ERK, MAPK, FIBROBLAST GROWTH FACTOR RECEPTOR, ETV5, TET2, EMBRYONIC STEM CELLS

## Abstract

Dynamic equilibrium of extracellular signal-regulated kinase (ERK) activity is regulated elaborately by multiple feedback loops to ensure the normal self-renewal of mouse embryonic stem cells (mESCs). Previous studies on mESCs have demonstrated that the negative feedback loops are engaged to prevent the overactivated ERK phosphorylation (pERK). It is not clear whether there is any positive feedback loop involved to maintain a minimum of pERK in mESCs. Here, we found that blocking fibroblast growth factor (FGF)-ERK pathway by chemical PD0325901 downregulated the transcription of E26 transformation-specific (ETS) family transcription factor Etv5 in mESCs. In turn, knockout (KO) of Etv5 by CRISPR/Cas9 decreased pERK. Moreover, Etv5 KO enhanced the DNA methylation at promoter of fibroblast growth factor receptor 2 (Fgfr2) by downregulating DNA hydroxylase Tet2, which further decreased the expression of Fgfr2 in mESCs. Collectively, a positive feedback loop of regulating pERK was revealed in mESCs, which was mediated by Etv5-Tet2-Fgfr2 axis. Our findings provide a new paradigm for pERK regulation in mESCs and will be useful to understand the cell fate determination during early embryo development.

## INTRODUCTION

Pluripotency is established and maintained by the cooperation of transcription factors (TFs), epigenetic factors and signaling pathways. A handful of TFs have been found to play pivotal roles in pluripotency maintenance over the past decade (Niwa, 2018). A core trinity of those TFs is Oct4, Sox2, and Nanog (OSN). Genetic loss-of function studies identified genes that directly or indirectly interacted with OSN are also essential for pluripotency, which include Sall4, Foxd3, Esrrb, Tbx3, Nr5a2, Dax1, Prdm14, Stat3, Myc family members, and Krüppel-like family members (Loh et al., 2015). Although considerable progress has been made on TFs network of pluripotency, the picture of pluripotency regulation is still not fully understood.

The E26 transformation-specific (ETS) genes constitute a large family of TFs and can be divided into five subfamilies, including ETS, ERG, ELG, TEL, and PEA3. The PEA3 subfamily is composed of Etv1, Etv4 and Etv5, which share a conserved ETS domain and two transactivation domains. Etv5, also called ets-related molecule (ERM), functions essentially in kidney development, lung development, and reproduction (Findlay et al., 2013). Etv5 is also an important member of TFs network anchored on OSN in mouse embryonic stem cells (mESCs) (Zhou et al., 2007). However, the exact roles of Etv5 in pluripotency regulation have been elusive for a long time until recently. Etv5, as well as Nr0b1, was proposed as the early regulator of somatic reprogramming (Lujan et al., 2015). We previously found that Etv5 could remarkably promote the efficiency of generating induced pluripotent stem cells (iPSCs) when combined with Yamanaka factors and facilitate mesenchymal–epithelial transition (MET) through Tet2-miR200s-Zeb1 regulation axis (Zhang et al., 2018a).

Unexpectedly, Etv4 and Etv5 appeared to be dispensable for maintenance of mESCs undifferentiated state. Etv4 and Etv5 double knockout (Etv4/5 dKO) did not affect the expression of Oct4 and Nanog in mESCs. However, Etv4/5 dKO decreased the proliferation of mESCs and impaired ectoderm differentiation (Akagi et al., 2015). By contrast, our previous study revealed that Etv5 knockdown (KD) could decrease the expression level of Tet2 and 5-hydroxymethylcytosine (5hmC) in mESCs and delayed the primitive endoderm (PrE) differentiation by downregulating Gata6 (Zhang et al., 2018a). These findings suggest Etv5 may be engaged in the initiation of differentiation. However, the mechanism of initiating differentiation by Etv5 remains largely unknown.

Fibroblast growth factors (FGFs) and their receptor tyrosine kinases play critical roles in the early embryo development and pluripotency regulation. The FGF signaling pathway is activated by a ligand-receptor interaction which leads to the autophosphorylation of tyrosine residues in the intracellular region of a FGF receptor (FGFR). Then the signal further activates four distinct pathways, including Janus kinase/signal transducer and activator of transcription (Jak/Stat) pathway, phosphoinositide phospholipase C pathway, phosphatidylinositol 3-kinase pathway, and mitogen-activated protein kinase/extracellular signal-regulated kinase (MAPK/ERK) pathway (Lanner and Rossant, 2010). Much evidence has demonstrated that autocrine FGF4/ERK signaling is needed for mESCs to exit from self-renewal and initiate differentiation (Kunath et al., 2007). Tracing the expression of Fgfr1 and Fgfr2 in early mouse embryos found that both Fgfr1 and Fgfr2 were required to mediate FGF signaling during PrE development. In addition, A permissive function of FGF4/FGFR1 signaling was found to mediate exit from pluripotency (Kang et al., 2017; Molotkov et al., 2017). However, the relationship between FGFRs and pluripotent TFs in mESCs is still unclear.

In this study, we investigated the interaction between Etv5 and FGF-ERK pathway in mESCs and found that blocking FGF-ERK pathway by MEK inhibitor PD0325901 downregulated the transcription of Etv5. By contrast, Etv5 KO reduced the pERK in mESCs. Mechanistically, Etv5 KO enhanced the DNA methylation of Fgfr2 promoter by downregulate Tet2, which further reduced the expression of Fgfr2 in mESCs. Briefly, our findings revealed a positive feedback loop of regulating pERK in mESCs, which was mediated by Etv5-Tet2-Fgfr2 axis. Interaction between Etv5 and FGF/ERK pathway found in this study will provide insights to understand the initiation of mESCs differentiation in vitro and lineage specification of embryo development in vivo.

## RESULTS

### Etv5 is a downstream target of FGF/ERK signaling pathway in mESCs

ETS transcription factors are often transcriptionally induced by FGF/ERK signaling during early embryo development (Lunn et al., 2007; Miya and Nishida, 2003; Willardsen et al., 2014). We asked whether Etv5, one member of the ETS transcription factors, was a downstream target of FGF/ERK signaling pathway in mESCs as seen in early embryo (Fig.1A). As mESCs were treated with MEK inhibitor PD0325901 with different doses (0, 0.5, 1.0, 1.5, 2.0, 2.5 μM), the pERK was expectedly downregulated in mESCs as the dose of PD0325901 increased (Fig.1B). The expression of Etv5 mRNA in mESCs was also decreased as the dose of PD0325901 increased (Fig.1C). These results indicate the transcriptional expression of Etv5 is positively regulated by pERK in mESCs. Furthermore, we investigated the time course change of pERK and Etv5 mRNA expression in mESCs which were treated with 1 μM PD0325901 for different hours (0, 3, 6, 9, 12h). The pERK was decreased more dramatically as the mESCs were treated with PD0325901 for a longer time (Fig.1D). The expression of Etv5 mRNA was correspondingly reduced as the pERK decreased in mESCs at distinct time point (Fig.1E). Together, these results indicate that the transcription of Etv5 is activated by FGF/ERK signaling pathway in mESCs.

**Fig.1.**
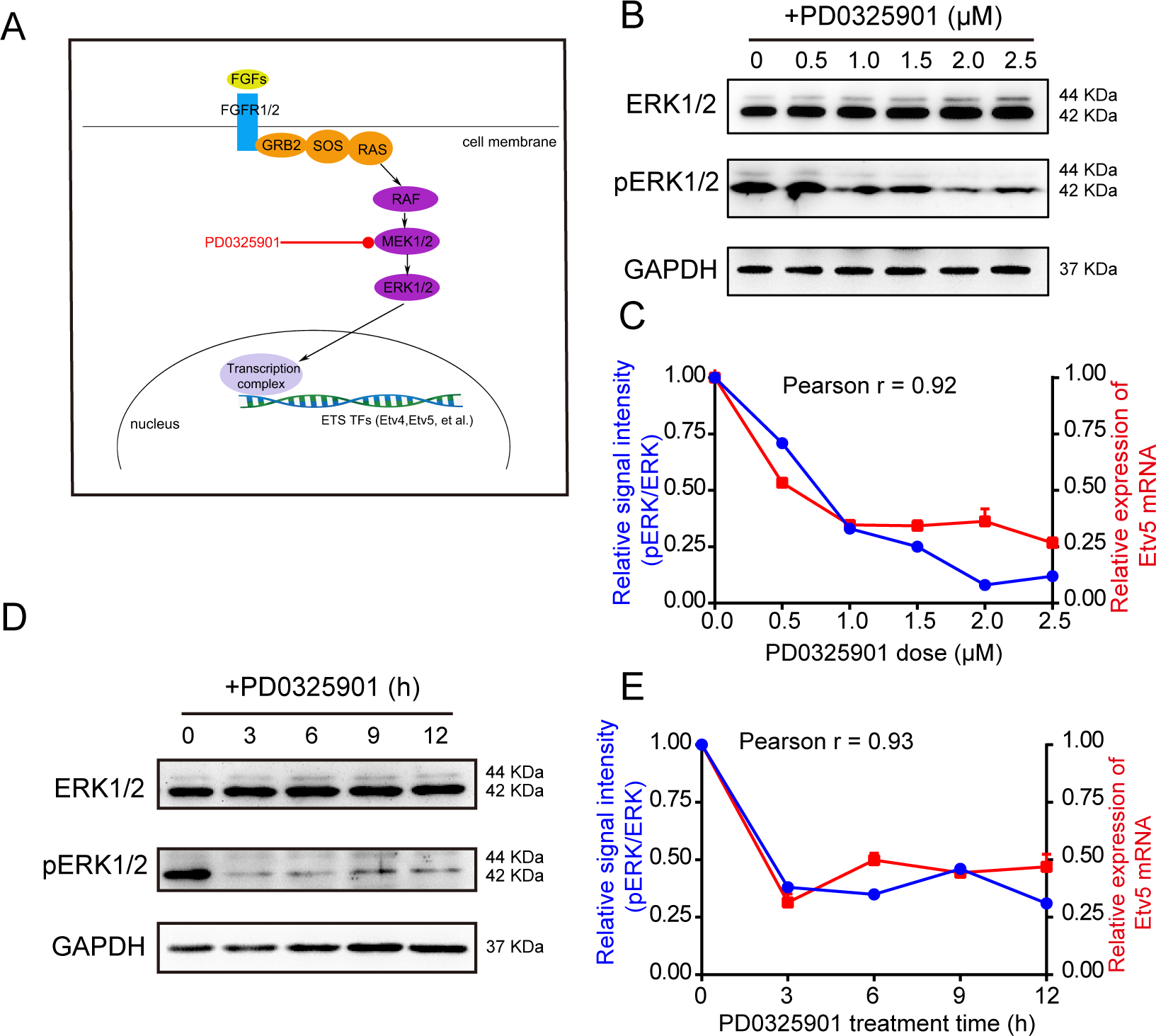
Etv5 is regulated by FGF/ERK pathway in mESCs. (A) Illustration of FGF/ERK pathway and transcriptional regulation of E26 transformation-specific (ETS) family members (Etv4 and Etv5). Small molecule chemical PD0325901 is a MEK inhibitor that blocks pERK. (B) Western blot of pan ERK and pERK in mESCs treated by different doses of PD0325901. GAPDH is used as internal control. (C) Relative pERK/pan ERK signal intensity (left Y-axis) and relative Etv5 mRNA expression (right Y-axis) in mESCs treated by different doses of PD0325901. Pearson’s correlation coefficient r is 0.92. (D) Western blot of pan ERK and phosphorylated ERK (pERK) in mESCs treated with PD0325901 (1μM) for different time points. GAPDH is used as internal control. (E) Relative pERK/pan ERK signal intensity (left y-axis) and relative Etv5 mRNA expression (right y-axis) in mESCs treated with PD0325901 (1μM) for different time points. Pearson’s correlation coefficient r is 0.93.

### Decreased pERK in Etv5 KO mESCs

Then we asked whether Etv5 had a feedback regulation of FGF/ERK pathway in mESCs. Etv5 KO mESCs were generated by CRISPR/Cas9 (Table S1). The genomic sequence located in the exon 7 of Etv5 gene was deleted and validated by Sanger sequencing (Fig.2A). Three Etv5 KO mESCs clones (B10, B13, and B23) were used for further investigation. Surprisingly, we found that pERK was all decreased in the Etv5 KO mESCs clones (B10, B13, and B23) when compared with wild type (WT) cell line J1. By contrast, Etv5 KO had no effect on the expression level of pan ERK (Fig.2B). The pERK level of KO versus WT ranged from 13% to 70% (Fig.2C). Collectively, these results suggest that Etv5 can regulate FGF/ERK pathway in mESCs with a positive feedback.

**Fig.2.**
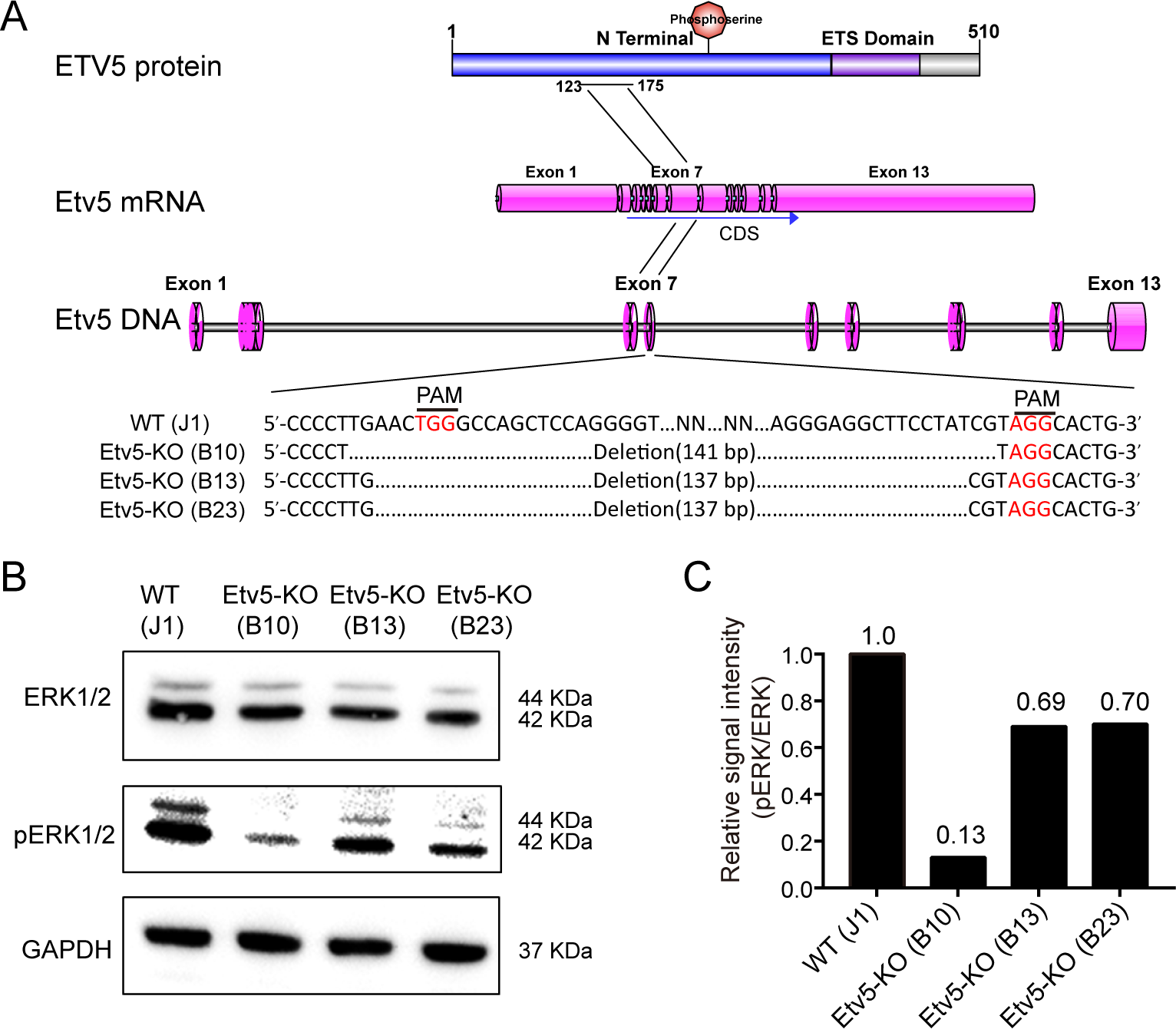
Etv5 KO decreases pERK in mESCs. (A) Illustration of Etv5 KO by CRISPR-Cas9. The position of a deletion in Etv5 DNA, mRNA, and protein is shown (top). Sanger sequencing of three representative KO clones (B10, B13, and B23) and wild type (WT) J1 mESC is also shown (bottom). (B) Western blot of pan ERK and pERK in WT J1 mESCs and Etv5 KO clones. GAPDH is used as internal control. (C) Relative pERK/pan ERK signal intensity in WT J1 mESCs and Etv5 KO clones.

### Phenotype relavant to decreased pERK in Etv5 KO mESCs

To study the impact of decreased pERK in Etv5 KO mESCs, we further investigated the phenotypes on proliferation, differentiation, and apoptosis.

FGF/ERK pathway is involved in stimulating cellular proliferation (Coutu and Galipeau, 2011). So we asked whether decreased pERK caused by Etv5 KO could slow down the proliferation of mESCs. Growth curve assay showed that Etv5 KO mESCs clones proliferated at a slower rate when compared to WT mESCs (Fig.3A). The proliferation defect of Etv5 KO mESCs is in line with our previous observation on Etv5 KD mESCs (Zhang et al., 2018a).

**Fig.3.**
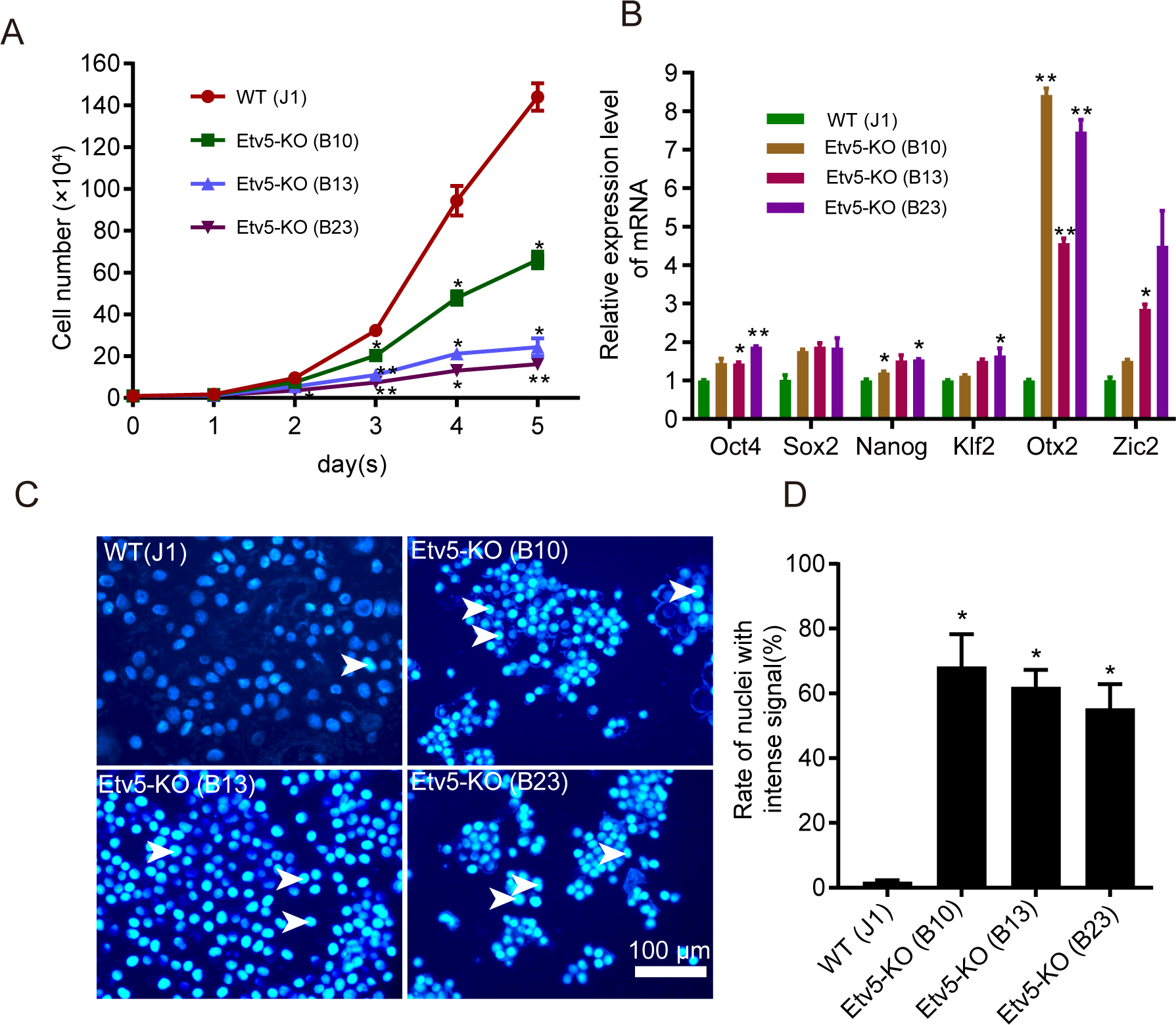
Phenotype of Etv5 KO mESCs. (A) Growth curve of WT J1 mESCs and Etv5 KO clones. \**P*<0.05, \*\**P*<0.01 (n=3, One-way ANOVA with Dunnett’s test). (B) Relative mRNA expression level of pluripotency-relevant genes in WT J1 mESCs and Etv5 KO clones. \**P*<0.05, \*\**P*<0.01 (n=3, One-way ANOVA with Dunnett’s test). (C) Hoechst 33258 staining of WT J1 mESCs and Etv5 KO clones. Arrowheads, cells with intense signal of Hoechst 33258 staining. Scale bar, 100 μm. (D) Rate of cells with intense signal of Hoechst 33258 staining in the population of WT J1 mESCs and Etv5 KO clones. \**P*<0.05 (n=3, One-way ANOVA with Dunnett’s test).

FGF/ERK is suggested to trigger lineage commitment from pluripotent state in mESCs (Kalkan and Smith, 2014). So we tested whether decreased pERK caused by Etv5 KO could enhance the pluripotency. We compare the gene expression level of naïve (Nanog, Klf2) and primed (Otx2, Zic2) pluripotency between Etv5 KO and WT mESCs. However, no considerable upregulation of Nanog and Klf2 was found between Etv5 KO and WT mESCs, although a slight increase was found in some Etv5 KO clones. Unexpectedly, Otx2 and Zic2 were upregulated sharply in Etv5 KO mESCs when compared to WT mESCs (Fig.3B). The other pluripotent properties, including alkaline phosphatase activity, expression of Oct4 and Sox2, were compared and no difference was found between Etv5 KO and WT mESCs (Fig.S1A-B). As Otx2 restricts mESCs to differentiate into primordial germ cell-like cells (PGCLCs) (Zhang et al., 2018b), the upregulation of Otx2 in Etv5 KO mESCs triggered us to examine whether Etv5 KO mESCs have a defect on PGCLCs differentiation. However, similar efficiency of PGCLCs differentiation was found between Etv5 KO and WT mESCs, indicating that Etv5 has little effect on the specification of PGC fate (Fig.S1C-D).

Genetic loss of ERK can increase apoptosis in mESCs (Chen et al., 2015). So we asked whether reduced pERK caused by Etv5 KO could also increase apoptosis in mESCs. The cells were stained with fluorescent dye Hoechst 33258 and intense signals were found in the nuclei of Etv5 KO mESCs when compared that with WT mESCs (Fig.3C-D). However, we observed no prominent cell death of Etv5 KO mESCs in our routine culture. So we further stained the cells with Annexin V and propidium iodide (PI). There is no significant difference when compared the rate of distinct cell populations (Annexin V^−^/PI^−^, Annexin V^−^/PI^−^, Annexin V^−^/PI^−^, and Annexin V^−^/PI^−^) in WT J1 mESCs with that in Etv5 KO clones (B10, B13, and B23) (Fig.S2A-B). Taken together, Etv5 KO can lead to chromatin condensation, which may make mESCs prone to apoptosis. (Fig.S2A-B)

### Etv5 maintains pERK by positively regulating Fgfr2 in mESCs

To dissect the molecular mechanism of reduced pERK caused by Etv5 KO in mESCs, we analyzed our previous RNA-seq data for Etv5 KD in mESCs (Zhang et al., 2018a) and investigated the expression change of genes in FGF/ERK pathway and genes which dephosphorylate pERK (Fig.4A). Interestingly, Fgfr2 and Fgfr3 were significantly downregulated in Etv5 KD mESCs when compared to nonsense control (NC) mESCs (FDR<0.001, |Fold change|>1.5) (Fig.4B, Table S2). Then we compared the expression abundance of Fgfr2 and Fgfr3 in mESCs and found that Fgfr2 was dominantly expressed in mESCs (Fig.4C). Therefore, we focused on Fgfr2 and further validated this finding in Etv5 KO mESCs by RT-qPCR (Fig.4D). The downregulated phosphorylation of Fgfr2 was also confirmed in Etv5 KO mESCs by Western blotting (Fig.4E-F). Briefly, these results indicate that Etv5 maintains pERK in mESCs by mainly regulating the expression level of Fgfr2 with a positive way.

**Fig.4.**
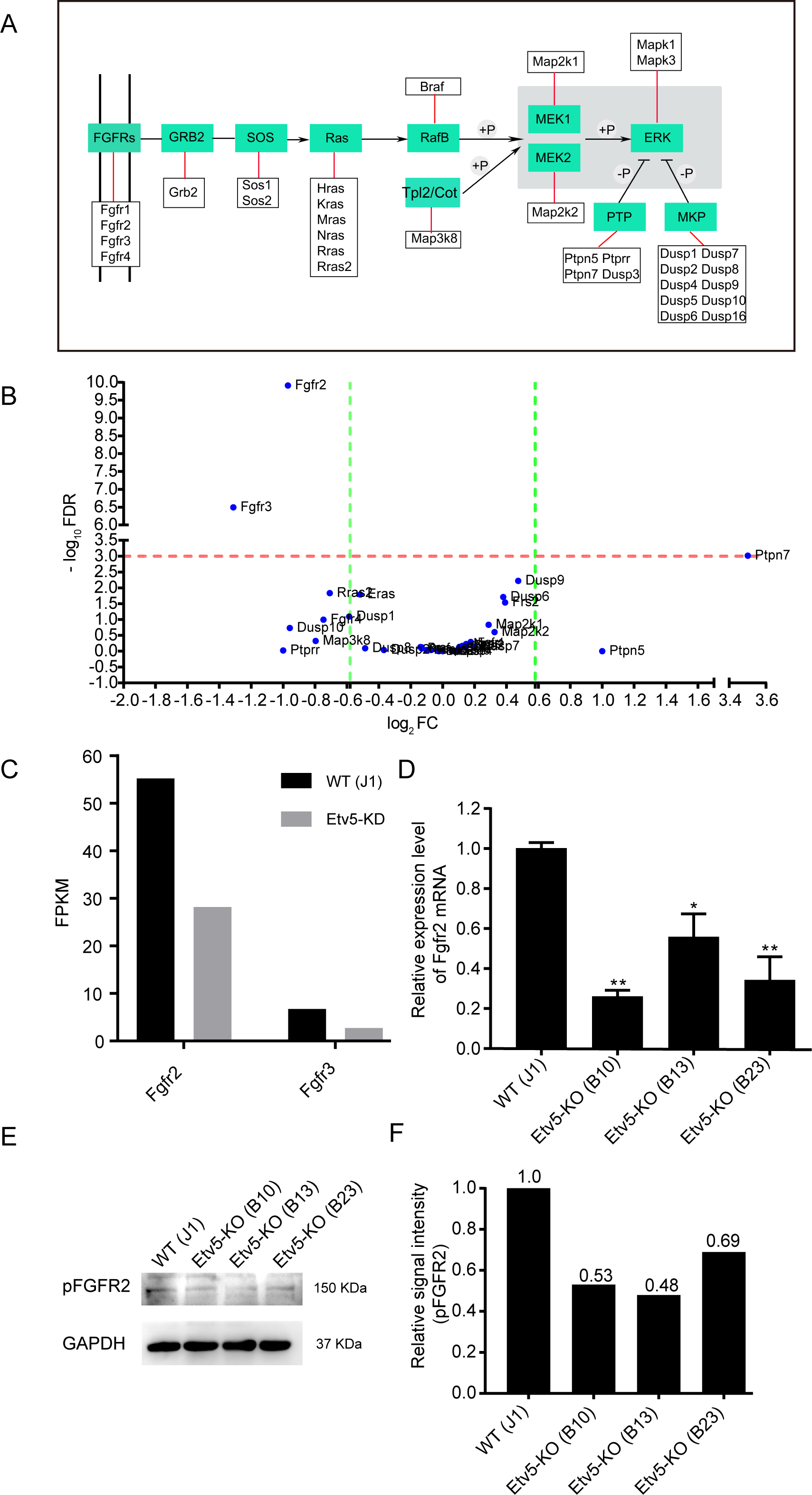
Etv5 KO decreases the expression level and pFgfr2 in mESCs. (A) Schematic diagram of regulating pERK by FGF-ERK pathway and phosphatase. +P, phosphorylate, -P, dephosphorylate. (B) Genes of FGF-ERK pathway and genes that dephosphorylate pERK expressed differentially between Etv5 KD and NC mESCs. Red dotted line, false discovery rate (FDR) is 0.001. Green dotted line, absolute value of fold change (FC) is 1.5. (C) Fragments Per Kilobase Million (FPKM) value of Fgfr2 and Fgfr3 in Etv5 KD and NC mESCs. (D) Relative mRNA expression level of Fgfr2 in WT J1 mESCs and Etv5 KO clones. \**P*<0.05, \*\**P*<0.01 (n=3, One-way ANOVA with Dunnett’s test). (E) Western blot of pFGFR2 in WT J1 mESCs and Etv5 KO clones. GAPDH is used as internal control. (F) Relative pFGFR2 signal intensity in WT J1 mESCs and Etv5 KO clones.

### Etv5-Tet2-Fgfr2 regulation axis in mESCs

Furthermore, we asked how Etv5 regulated the transcription of Fgfr2 in mESCs. We firstly tested whether Etv5 had a direct activation on Fgfr2 promoter. To choose the potential region of Fgfr2 promoter, we analyzed the H3K4me3 ChIP-seq data of different mESC lines (E14, J1, and R1) and selected a region (chr7:130,264,821-130,266,989) with enriched peaks for luciferase assay (Fig.5A). Co-transfection of reporter plasmids and Etv5-expressing plasmids showed that Etv5 could generate about two-fold higher luciferase activity than control (Fig.5B). We analyzed the sequence of Fgfr2 promoter and found only one conserved motif (GGAAGT) for Etv5 binding. However, no obvious change of luciferase activity was observed when the conserved motif was mutated (Fig.5B).

**Fig.5.**
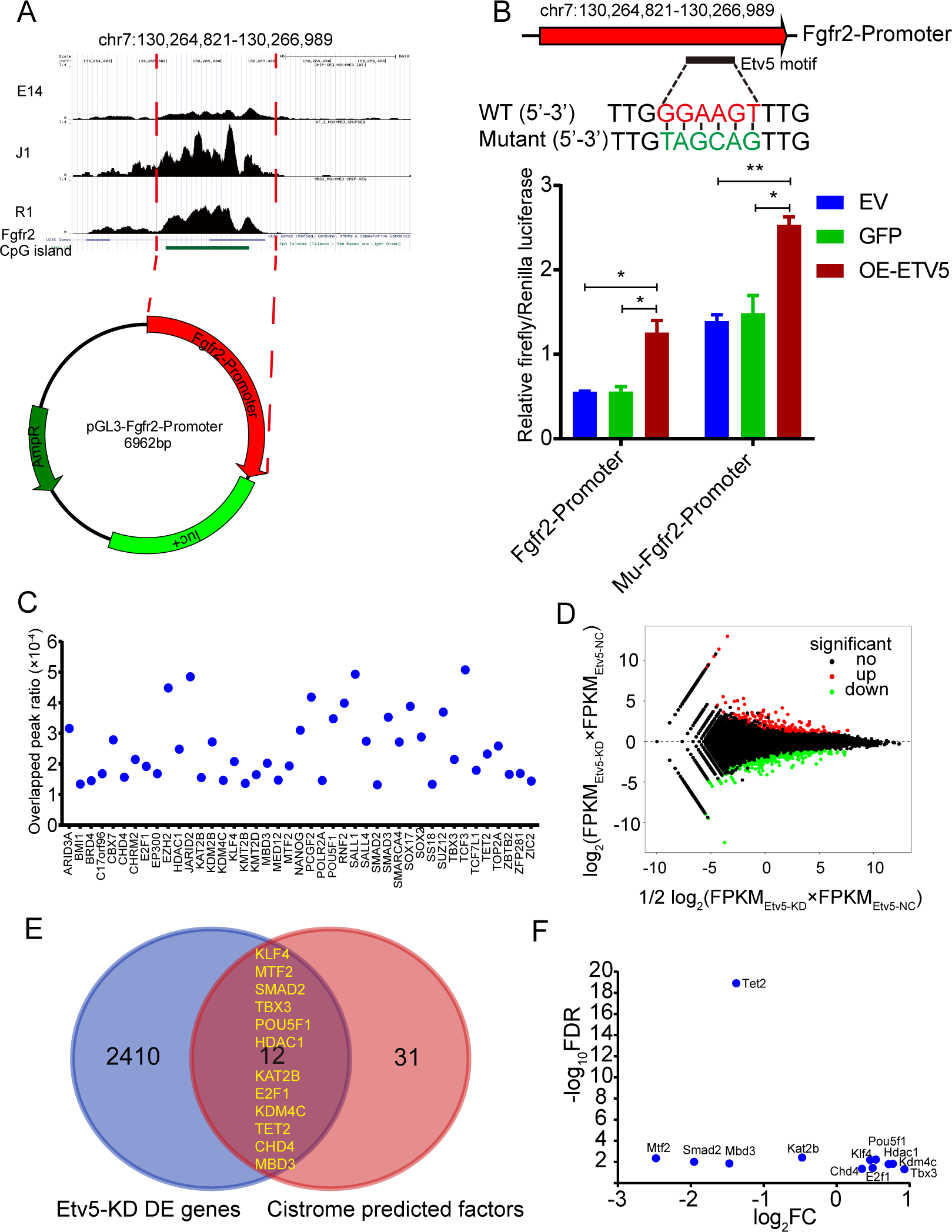
Etv5 regulates the expression of Fgfr2 potentially mediated by Tet2 in mESCs. (A) H3K4me3 ChIP-Seq of three representative mESC lines (E14, J1, and R1). UCSC genome browser snapshot showing the enriched H3K4me3 peaks at the putative promoter region of mouse Fgfr2 gene (chr7:130,264,821-130,266,989). The diagram of cloning putative Fgfr2 promoter into luciferase (Luc+) reporter plasmid pGL3 is also shown. Green bar, the CpG island region. (B) Luciferase assay of Fgfr2 promoter and Etv5-binding-motif-mutated Fgfr2 promoter. Empty vector (EV) is used as vehicle control. Vector expressing green fluorescent protein (GFP) is used as negative control. Vector expressing Etv5 is used to detect luciferase activity of Fgfr2 promoter and Etv5-binding-motif-mutated Fgfr2 promoter. *P<0.05, **P<0.01 (n=3, One-way ANOVA with Dunnett’s test). (C) Toolkit for Cistrome Data Browser showing the overlapped peak ratio of transcription factors and chromatin regulators binding to the putative promoter region of mouse Fgfr2 gene (chr7:130,264,821-130,266,989). (D) MA plot showing the genes differentially expressed between Etv5 KD and NC mESCs. Genes with significant change (p<0.05) in Etv5 KD mESCs versus NC mESCs are shown. Red dots, significantly upregulated genes in Etv5 KD mESCs versus NC mESCs. Green dots, significantly downregulated genes in Etv5 KD mESCs versus NC mESCs. Black dots, genes with no significant changes in Etv5 KD mESCs versus NC mESCs. (E) Venn diagram showing 12 overlapped genes between Cistrome predicted transcription factors/chromatin regulators binding to Fgfr2 promoter and genes with significant change (p<0.05) in Etv5 KD mESCs versus NC mESCs. (F) Scatter plot showing the 12 overlapped genes differentially expressed between Etv5 KD mESCs and NC mESCs. FDR, false discovery rate. FC, fold change.

The minor effect of directly activating Fgfr2 promoter by Etv5 triggered us to investigate whether Etv5 regulates Fgfr2 expression mediated by other factors. We analyzed the potential Fgfr2 promoter region (chr7:130,264,821-130,266,989) by using toolkit for Cistrome data browser and found 43 transcription factors or chromatin regulators binding to this region (Fig.5C). Then we asked if the 43 candidate genes were among the list of differentially expressed genes caused by Etv5 KD. The RNA-seq data showed that there were 2422 differentially expressed genes between Etv5 KD mESCs and NC mESCs (FDR<0.05) (Fig.5D, Table S2). Out of the 43 candidate genes, Venn analysis uncovered 12 genes were included among the list of differentially expressed genes caused by Etv5 KD, including Klf4, Mtf2, Smad2, Tbx3, Pou5f1, Hdac1, Kat2b, E2f1, Kdm4c, Tet2, Chd4, and Mbd3. These genes are believed to mediate the regulation of Fgfr2 expression by Etv5 (Fig.5E). Of note, Tet2 was the gene affected by Etv5 with great significance (FDR=1.25×10^−19^) (Fig.5F, Table S2). Together, the results above indicate that Etv5 may regulate the transcription of Fgfr2 mediated by Tet2 in mESCs.

### Increased promoter DNA methylation of Fgfr2 in Etv5 KO mESCs

Finally, we tested whether the reduced expression of Tet2 caused by Etv5 KO could increase the promoter DNA methylation of Fgfr2 in mESCs. The results of RT-qPCR and Western blotting validated that Tet2 was significantly downregulated in Etv5 KO mESCs when compared to WT mESCs (Fig.6A-C). Then we performed integration analysis of Tet2 and H3K4me3 ChIP-seq data. The WashU genome browser track showed that there were overlapped peaks across the Fgfr2 promoter (chr7:130,264,821-130,266,989) (Fig.6D). The region (chr7:130,266,142-130,266,551), a part of the CpG island, was chosen for bisulfite sequencing. Interestingly, we found that the DNA methylation level was increased in Etv5 KO mESCs (16.7%) when compared to WT mESCs (2.5%) from the investigated region (Fig.6E). Collectively, these results indicate that Etv5 can regulate the transcription of Fgfr2 by modifying its promoter DNA methylation level, which is mediated by Tet2.

**Fig.6.**
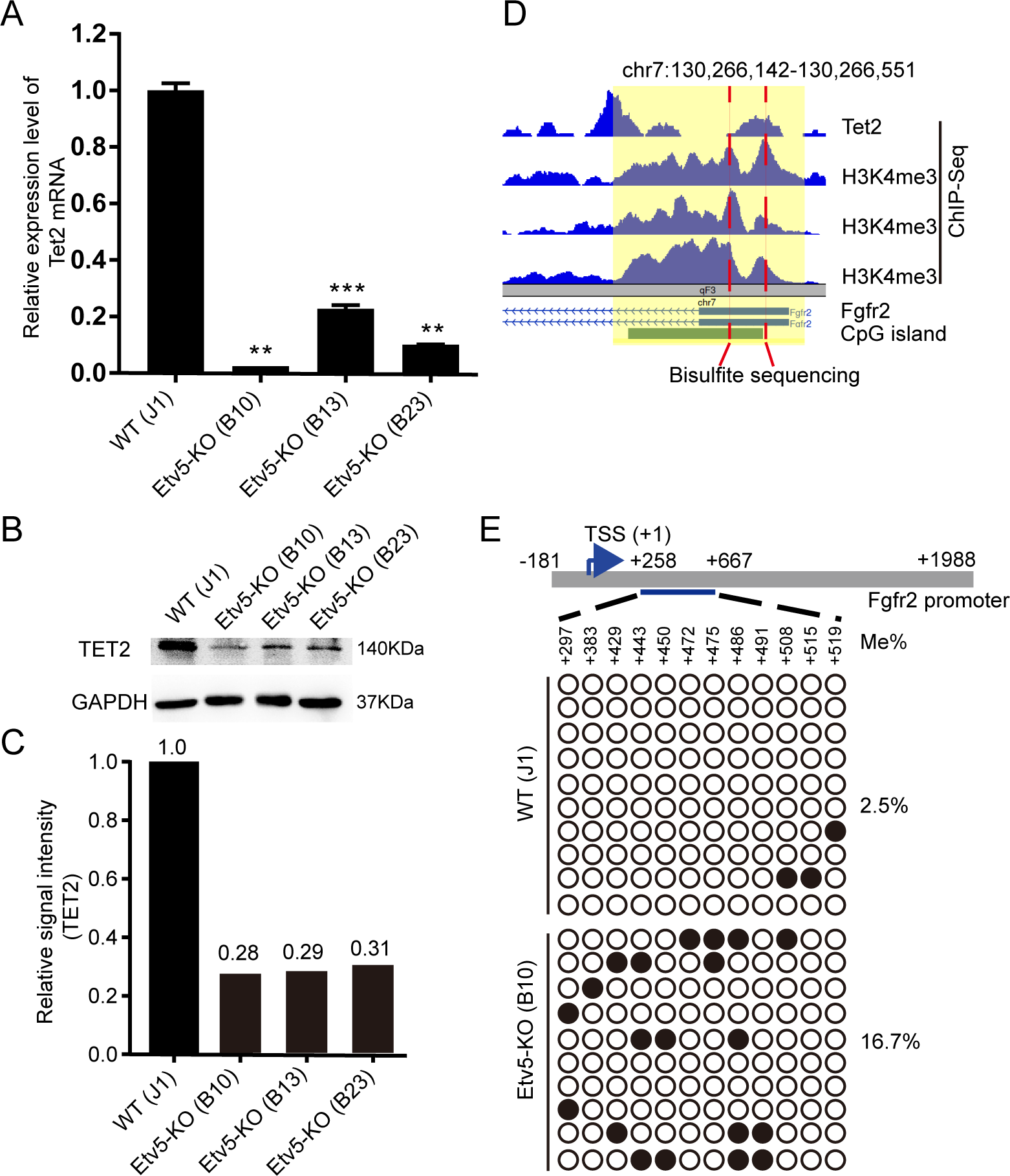
Increased DNA methylation at Fgfr2 promoter in Etv5 KO mESCs. (A) Relative mRNA expression level of Tet2 in WT J1 mESCs and Etv5 KO clones. \*\**P*<0.01, \*\*\**P*<0.001 (n=3, One-way ANOVA with Dunnett’s test). (B) Western blot of Tet2 in WT J1 mESCs and Etv5 KO clones. GAPDH is used as internal control. (C) Relative Tet2 signal intensity in WT J1 mESCs and Etv5 KO clones. (D) WashU genome browser snapshot showing the enriched Tet2 and H3K4me3 peaks at the putative promoter region of mouse Fgfr2 gene (chr7:130,264,821-130,266,989). A representative region (chr7:130,266,142-130,266,551) overlapped among Tet2 peaks, H3K4me3 peaks, and CpG island (green bar) is selected for bisulfite sequencing analysis. (E) Bisulfite sequencing of Fgfr2 promoter for WT J1 mESCs and Etv5 KO clone (B10). Open and filled circles indicate unmethylated and methylated CpG dinucleotides, respectively. The differentially methylated sites relative to transcription start site (TSS) of Fgfr2 are shown. The percentage of methylated sites (Me %) is also presented.

## DISCUSSION

In this study, a positive loop of regulating pERK mediated by Etv5 was found in mESCs. Blocking FGF-ERK pathway could downregulate the transcription of Etv5 in mESCs, while Etv5 KO could decrease the pERK in mESCs through Etv5-Tet2-Fgfr2 regulation axis (Fig.7A-B).

**Fig.7.**
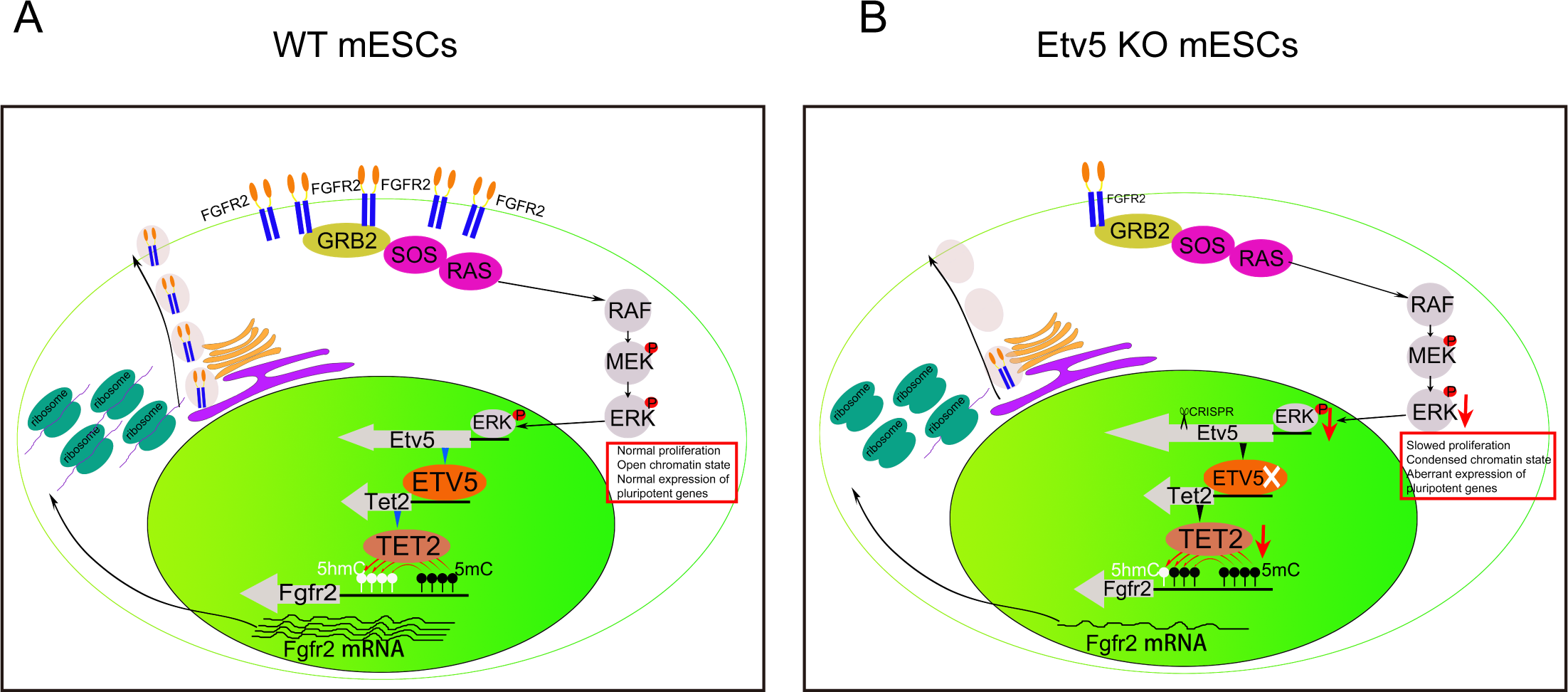
Positive feedback loop model of regulating pERK in mESCs mediated by Etv5-Tet2-Fgfr2 axis. (A) In WT mESCs, the active FGF-ERK pathway can activate the transcription of Etv5. In return, Etv5 can promote the expression of Fgfr2, which is mediated by Tet2. (B) In Etv5 KO mESCs, Loss of Etv5 leads to the downregulation of Tet2, which further increases the methylation level of Fgfr2 promoter and results in the downregulation of Fgfr2. The pERK is accordingly decreased.

Activation of ERK can increase the transcriptional output via release of paused RNA polymerase (Williams et al., 2015). Transcriptional regulation of ETS family members by FGF-ERK pathway seems well conserved among early embryos, germline stem cells, and ESCs. Transcription of Etv5, as well as Pea3, is tightly linked with active FGF signaling during gastrulation and somitogenesis stages of zebrafish embryos (Znosko et al., 2010). A similar finding was discovered during mouse epiblast specification. The expression of Etv5, as well as Etv4, is controlled by FGFR1 signaling within the epiblast cells at E3.5 (Kang et al., 2017). Etv5 is also essential for maintaining the self-renewal of spermatogonial stem cells. The expression of Etv5 can be induced by the activated FGF2-ERK pathway. However, excessive stimulation of FGF2-ERK pathway can generate an abundance of Etv5, which upregulates Bcl6b expression and results in germ cell tumor (Ishii et al., 2012). In this study, we provide extra evidence that transcriptional regulation of Etv5 by FGF2-ERK pathway is also conserved in mESCs. This finding represents a typical crosstalk between extracellular FGF-ERK signaling pathway and transcriptional networks in mESCs.

The different pERK level has diverse roles in mESCs. Although chemical inhibition of pERK is beneficial for the self-renewal of mESCs (Ying et al., 2008), the minimum activity of ERK signaling is essentially required for proliferation, anti-apoptosis, and genomic stability of mESCs. By contrast, over-activated ERK signaling can impair the self-renewal of mESCs and activate developmental genes (Ma et al., 2016). Etv5 KO resulted in lower pERK level than the minimum for normal mESCs maintenance. Our finding suggests that Etv5 may be indispensable to maintain the proliferation, anti-apoptosis, and genomic stability of mESCs. Consistent with this prediction, the Etv5 KO mESCs exhibited slower proliferation and a tendency to apoptosis. Our results refute the view that Etv5 is not essential for mESCs’ maintenance(Chen et al., 2008).

There are elaborate feedback systems to keep the minimum activity of ERK signaling in mESCs. On the one hand, negative feedback would be activated once the pERK level was higher than the minimum. Cellular strategies of negative feedback regulation include fast-acting mechanisms and slow-acting mechanisms (Lake et al., 2016). The fast-acting mechanisms utilize the downstream components of the cascade (such as ERK1/2, RSK1, and RSK2) to directly phosphorylate various upstream components and downregulate the activity of ERK signaling (Nett et al., 2018; Sturm et al., 2010). The slow-acting mechanisms utilize the de novo expression of proteins (such as DUSPs and Sprouty proteins) to downregulate the activity of ERK at multiple nodes in the pathway with cellular context dependence (Li et al., 2007; Yu et al., 2011). On the other hand, positive feedback is expected to be activated when the pERK level was lower than the minimum. Experimental evidence has demonstrated that fast-acting positive feedback loop can be launched by ERK via inactivation of the inhibitor RKIP (Shin et al., 2009). However, so far there has been little evidence to prove the existence of slow-acting positive feedback loop for regulating ERK activity. In this study, we provide the first evidence that de novo expression of Etv5 can be induced by ERK and in turn Etv5 can upregulate the activity of ERK by increasing the expression abundance of Fgfr2 in mESCs (Fig.7A). Our finding represents a slow-acting positive feedback loop of regulating the ERK signaling pathway.

Fgf receptors (Fgfr1 and Fgfr2) are involved in the specification of PrE and epiblast within the inner cell mass (ICM) of the mouse blastocyst. High expression level of Fgfr1 ranges from the early blastocyst (E3.25) to late blastocyst (E4.5), suggesting Fgfr1 is a pan-ICM receptor. By contrast, high expression level of Fgfr2 is only observed from mid blastocyst (E3.5) to late blastocyst (E4.5), suggesting Fgfr2 is a PrE-biased receptor (Kang et al., 2017). The molecular mechanism of initiating Fgfr2 is unclear. Etv5, as well as Etv4, was indicated to regulate FGF-ERK signaling pathway as negative feedback at early blastocyst (Kang et al., 2017). Our findings in mESCs refuted this view and suggested that Etv5 could regulate FGF-ERK signaling pathway as positive feedback by promoting the expression of Fgfr2. Our findings also provided the clues to understand the possible roles of Etv5 and Etv4 at early blastocyst. The uncommitted ICM cells at early blastocyst express only FGFR1 to receive FGF4 signal and exhibit low ERK activity, which is supposed to trigger the expression of Etv4 and Etv5. Subsequently, Etv4 and Etv5 may initiate the expression of Fgfr2 in ICM cells in a salt-and-pepper pattern at mid blastocyst. These ICM cells expressing both Fgfr1 and Fgfr2 could receive more FGF4 signal and exhibit high ERK activity, which further promote the specification of PrE fate at the late blastocyst (Kang et al., 2017; Kang et al., 2013).

Our previous study has indicated that Etv5 is essentially needed for PrE differentiation, which may be mediated by Tet2 (Zhang et al., 2018a). In this study, we further demonstrated that Etv5 could regulate the DNA methylation of Fgfr2 promoter mediated by Tet2, which further impacted on ERK activity in mESCs. This finding clarified the molecular mechanism of regulating PrE differentiation by Etv5. However, unlike TET1 and TET3, TET2 does not contain a DNA-binding domain (Rasmussen and Helin, 2016), suggesting that TET2-mediated oxidization of 5-methylcytosine (5mC) into 5-hydroxymethylcytosine (5hmC) at Fgfr2 promoter should be helped by additional recruiters. Whether Etv5 could recruit Tet2 to at Fgfr2 promoter needs to be investigated in the future.

In sum, our study offers a new paradigm of regulating ERK signaling pathway in mESCs through slow-acting positive feedback loop, which is mediated by Etv5-Tet2-Fgfr2 axis (Fig.7A-B). FGF-ERK signaling pathway plays fundamental roles in proliferation, differentiation, apoptosis, migration, and genome stability during development (Ornitz and Itoh, 2015). However, aberrant activation of FGF-ERK signaling pathway often leads to tumorigenesis or carcinogenesis (Liu et al., 2018). Our findings in this study will provide useful clues to understand the cell fate determination during early embryo development and explore new strategies for cancer treatment.

## MATERIALS AND METHODS

### Cell culture

The J1 mESCs (ATCC) were routinely cultured on mitotically inactivated mouse embryonic fibroblast feeder layer with serum-containing mESCs medium. The serum-containing mESCs medium was composed of high glucose DMEM (HyClone) supplemented with 1000 U/mL recombinant mouse leukaemia inhibitory factor (LIF) (Millipore), 100 μM non-essential amino acids (NEAA) (Gibco), 1 mM L-glutamine (Gibco), 100 U/ml penicillin and 100 μg/ml streptomycin (HyClone), 100 μM β-mercaptoethanol (Sigma), and 15% FBS (Gibco). For CRISPR/Cas9 KO experiment, the J1 mESCs were grown on 0.1% gelatin-coated tissue culture plates with modified serum-containing mESCs medium as we previously described (Zhang et al., 2018a). The differentiation of PGCLCs from J1 mESCs and Etv5 KO mESCs was carried out according to a protocol we recently reported (Li et al., 2019).

### Generation of Etv5 KO mESCs by CRISPR/Cas9

A double sgRNA was designed to target the exon 7 of Etv5 and cloned into the PX459M (Miaolingbio). Then the designed CRISPR/Cas9 vectors were transfected into J1 mESCs using Lipofectamine 3000 (Invitrogen). After 24 hours, the transfected mESCs were treated with 1.5 μg/ml puromycin (Solarbio) for 48 hours and re-plated for single colony isolation. Individual colonies were picked up after five days’ expansion and screened by genotyping PCR. KO colonies were further validated by Sanger sequencing. The sgRNAs’s oligonucleotide sequences and primers for genotyping PCR are listed in Table S3.

### Western blot

The J1 mESCs and Etv5 KO mESCs were collected and lysed in RIPA buffer (Solarbio) with 1×protease inhibitor cocktail (Roche). When phosphorylated proteins were detected, additional 1×PhosSTOP (Roche) was added into the lysis buffer. The extracted proteins were diluted and boiled 5 minutes for SDS-PAGE. Then the proteins were transferred onto a polyvinylidene fluoride (PVDF) membrane (Millipore) using a Mini Trans Blot system (Bio-Rad). The semi-dry transfer condition of TET2 and pFGFR2 is 15V, 1.5 hours. Semi-dry transfer of other proteins in this study is 15V, 1 hour. The membranes were next blocked in TBST buffer (10 mM Tris-HCl pH 8.0, 150 mM NaCl and 0.5% vol/vol Tween-20) containing 5% skim milk powder for 1 hour at room temperature. After that, membranes were incubated with primary antibody at 4°C overnight. The next day, membranes were washed and then incubated with HRP-conjugated secondary antibody at room temperature for 1 hour. Signals were detected by enhanced chemiluminescence (Tanon) according to the manufacturer’s instructions. The antibodies used for western blot in this study are given in Table S4.

### RT-qPCR analysis

The total RNA of J1 mESCs and Etv5 KO mESCs was extracted with RNAsimple Total RNA Kit (TIANGEN) according to the manufacturer’s instructions. Then, the total RNA was converted to cDNA using FastKing RT Kit (TIANGEN). For qPCR, the cDNA was amplified using a Real-Time PCR System (Bio-Red) with BioEasy Master Mix (BIOER). The primers of RT-qPCR used in this study are listed in Table S5.

### Luciferase reporter assay

Fragments of putative Fgfr2 promoter and Etv5-motif-mutated Fgfr2 promoter were cloned into pGL3-basic reporter vector (Promega). The cloning primers are included in Table S6. For the luciferase assay, the reporter vectors were co-transfected into HEK293T cells with pRL-TK (Promega) and Etv5 over-expression plasmid pTrip-CAGG-Etv5 using Dual Luciferase Reporter Gene Assay Kit (Beyotime). Firefly luciferase activity and Renilla luciferase activity were measured 48 hours after transfection by using Microplate luminometer (Hamamatsu). The ratio of Firefly luciferase activity/Renilla luciferase activity was calculated and compared.

### Flow cytometry analysis

The PGCLCs (5×10^5^ cells) differentiated from WT and Etv5 KO mESCs were trypsinized and collected for flow cytometry analysis. The cells were resuspended in 30 μl PBS supplemented with Alexa Fluor 647 Mouse anti-SSEA1 (1:200, BD Biosciences) and FITC anti-mouse/rat CD61 (1:500, Biolegend) and incubated at 4°C for 30 minutes. Then the cells were washed twice in 1 ml PBS with 10% FBS and examined on a BD FACSAria™ III cell sorter (BD Biosciences).

### Alkaline phosphatase staining

The WT and Etv5 KO mESCs were fixed with 4% paraformaldehyde (Sigma) at room temperature for 10 minutes. Then the cells were washed with PBS three times and incubated with alkaline phosphatase staining solution as we previously described (Zhang et al., 2017). The images were finally taken and recorded under microscope (Nikon).

### Apoptosis relevant staining

For investigation of chromosome condensation, WT and Etv5 KO mESCs cultured in 6-well plate were washed with PBS three times and incubated with Hoechst33258 (1 ml/well, Wanleibio) for 20 minutes in the incubator (37°C, 5% CO_2_). After that, the cells were washed with PBS three times and images were taken and recorded with EVOS FL Imaging System (ThermoFisher Scientific).

For investigation of the early apoptotic markers, WT and Etv5 KO mESCs (~2×10^5^ cells) were collected and resuspended with 1×Binding Buffer containing Annexin V-FITC/PI (Sigma). The cells were incubated at room temperature in the dark for 5 minutes. Then the cells were collected by centrifugation and resuspended in 1×Binding Buffer for glass slide smear. The images were finally taken and recorded with EVOS FL Imaging System (ThermoFisher Scientific).

### Bisulfite Sequencing

Sodium bisulfite modification of genomic DNA (500 ng) of WT and Etv5 KO mESCs was performed according to the instruction of EZ DNA Methylation-Direct^TM^ Kit (Zymo Research). Methylation specific PCR for a fragment of Fgfr2 promoter was carried out as we previously described (Zhang et al., 2018a). The primers used for methylation specific PCR are listed in Table S6. Final PCR products were sequenced and analyzed with BiQ Analyzer.

### Immunofluorescence

WT and Etv5 KO mESCs were fixed with 4% paraformaldehyde (Sigma) for 20 minutes at room temperature. Then the cells were blocked in PBS containing 0.5% Triton X-100 and 0.1% bovine serum albumin (BSA) for one hour. After that, the cells were incubated with primary antibodies at 4°C overnight and followed by incubating with fluorophore-conjugated secondary antibody for one hour at room temperature. The antibodies used for immunofluorescence are listed in Table S4. Finally the cells were stained with DAPI (ThermoFisher Scientific) and images were taken by EVOS FL Imaging System (ThermoFisher Scientific).

### Cell proliferation assay

WT and Etv5 KO mESCs with the same cell number (1×10^4^ cells) were initially seeded into 24-well plates and thereafter counted every day. A total of five days were recorded. The growth curve was drawn and compared.

### Bioinformatics analysis

The RNA-seq data of Etv5 KD mESCs versus WT mESCs (SRA accession number, SRP111429) were used for bioinformatics analysis in this study. Briefly, the filtered reads were mapped the mouse reference genome (mm10) by TopHat2 (Kim et al., 2013)and Bowtie2(Langmead and Salzberg, 2012). Reconstruction of transcripts was carried out with Cufflinks (Trapnell et al., 2010). Transcripts abundance was quantified by software RSEM (Li and Dewey, 2011). The edgeR package (http://www.r-project.org/) was used for identifying differentially expressed transcripts across groups (Robinson et al., 2010). MA plot was drawn by using the OmicShare tools (www.omicshare.com/tools), a free online platform for data analysis. Venn analysis was performed by using Venn Diagrams (http://bioinformatics.psb.ugent.be/webtools/Venn/). mESCs ChIP-Seq data of H3K4me3 (GSM1385046, GSM2165937, GSM2052271) and TET2 (GSM2065691) were used for integration analysis. The toolkit for Cistrome Data Browser was used for predicting transcription factors and chromatin regulators which bind to a specified genomic interval (Zheng et al., 2019). UCSC Browser and WashU Browser were used to show peaks enrichment at a given genomic locus.

### Statistical analyses

One-way analysis of variance (ANOVA) with Dunnett’s multiple comparisons test was used in this study. Software GraphPad Prism 7.0 was used for the statistical analysis, and data are presented as means ± SD. \**P*<0.05, \*\**P*<0.01, \*\*\**P*<0.001.

## Acknowledgements

We thank Dr. Jinglong Zhang for helping on generation of Etv5 KO mESCs, Xiaomin Du for helping on bisulfite sequencing experiment, Mengfei Zhang for helping on apotosis staining experiment; Juqing Zhang and Zhenshuo Zhu for helping on vector construction in this work.

## Author contributions

Conceptualization: Sq.Z; Methodology: C.F, K.Z, Y.L, Hx.C, My.Z, N.L; Software: C.F, K.Z; Validation: C.F; Formal analysis: C.F, Y.W, Jl.H, Hy.W; Investigation: C.F; Resources: K.Z, Y.L, Hx.C; Data curation: Sq.Z, C.F; Writing - original draft: Sq.Z, C.F; Writing - review & editing: Sq.Z, Jl.H, Hy.W; Visualization: C.F, Y.L, Y.W; Supervision: Sq.Z; Funding acquisition: Sq.Z.

